# Liquid-liquid phase separation and amyloid aggregation in the 14-3-3 protein family

**DOI:** 10.64898/2026.01.07.698245

**Authors:** Saulė Rapalytė, Viktorija Karalkevičiūtė, Ieva Baronaitė, Dominykas Veiveris, Vytautas Smirnovas, Mantas Žiaunys, Darius Šulskis

## Abstract

The 14-3-3 protein family, with over 1300 binding partners, is one of the largest regulators of protein-protein interactions (PPIs) in eukaryotic cells. They recognise and bind to phosphorylation-related motifs in their partner proteins, creating scaffolds to stabilise proteins and allowing or inhibiting kinases’ access to their targets. 14-3-3 consists of seven isoforms (β, γ, ε, ζ, η, θ and σ), which are expressed across all tissues. Given their important role in PPIs, their dysregulation can contribute to a variety of diseases, such as cancer or neurodegenerative disorders like Creutzfeldt-Jakob’s, Parkinson’s or Alzheimer’s diseases. Pathological effects can arise due to loss of function, aberrant interactions or protein aggregation. Notably, 14-3-3 aggregates have been detected in Lewy bodies or cerebrospinal fluid, mirroring the presence of amyloid proteins, such as α-synuclein (α-syn) or β-amyloid. Toxic amyloid aggregation signals the onset of neurodegeneration, which can occur through misfolding of proteins or liquid-liquid phase separation (LLPS) – a process during which proteins condense into membraneless organelles.

Unlike other amyloidogenic proteins, there is little information on the conditions under which the 14-3-3 protein family members undergo LLPS or their relationship with amyloid aggregation. To address this gap, we examined all isoforms of 14-3-3 *in vitro* and observed the formation of amyloid aggregates and phase-separated droplets. Our results revealed that the ε and θ isoforms form amyloid-like fibrils that can accelerate α-syn aggregation. Furthermore, molecular crowding conditions promoted phase separation and aggregation in most 14-3-3 proteins. Finally, 14-3-3s incorporated together with α-syn to generate heterotypical droplets.

## Introduction

The 14-3-3 protein family is one of the key interaction hubs that coordinate protein-protein interactions (PPIs) in mammals and plants [1]. They can create scaffolds that facilitate protein binding with their partners, or they can block and dissociate complexes. Most commonly, 14-3-3s recognise phosphorylation motifs and help kinases or phosphatases reach their targets [2]. 14-3-3 isoforms are evolutionarily conserved [1] and expressed in most tissues, but especially in the brain, where they can comprise up to 1% of soluble proteins [2]. Thus, they also interact with numerous neurodegeneration-linked proteins such as tau [3], α-synuclein (α-syn) [4] and coordinate their PPIs [5].

A key pathological feature of tau and α-synuclein is that they form insoluble protein aggregates known as amyloids [6], which are considered to be the central hallmarks of Alzheimer’s and Parkinson’s diseases, respectively [7], [8]. Amyloid plaques often contain not only these proteins, but also their interaction partners that co-aggregate together due to spatial proximity and affinity [9]. One of the early steps that promotes protein aggregation is liquid-liquid phase separation (LLPS), during which proteins condense into membraneless droplets, resulting in a local increase in protein concentration. LLPS can facilitate interactions between proteins, DNA, and RNA [10], [11], [12], and also accelerate the aggregation of proteins due to a crowded environment [13].

14-3-3s are found in Lewy bodies due to co-interactions with α-synuclein [14] and can mitigate α-synuclein aggregation [4] as well as reduce the propagation of amyloid fibrils [15]. Notably, a recent study showed that potential 14-3-3 amyloid structures also exist in tumor-derived cell lines [16]. Beyond amyloidogenesis, 14-3-3s have been suggested as potential regulators of LLPS. Among 1370 analysed binding partners, roughly half of them had a high propensity for phase separation [17]. Some examples include the microtubule-binding protein tau, where 14-3-3ζ prevents both aggregation and condensation of phosphorylated tau [18]. 14-3-3s also stabilise tumour suppressor factor p53 [19] and bind to SARS-CoV-2 nucleocapsid protein in low micromolar concentrations [20]. Both p53 [21] and nucleocapsid [22] are phase-separating proteins with critical roles in cancer or SARS-CoV-2 replication. Despite these numerous interactions with LLPS-forming proteins, it remains unknown whether 14-3-3 proteins themselves are capable of undergoing phase separation and forming amyloid fibrils.

Recently, our laboratory showed that 14-3-3ζ can form short amyloid fibrils [23]; interestingly, other isoforms share conserved sequence features that may predispose them to similar behaviour [24]. In this study, we systematically investigate the amyloidogenic and LLPS propensities of all seven 14-3-3 isoforms *in vitro*. We combined computational predictions of aggregation-prone regions and LLPS-associated domains with experimental analyses of protein stability, aggregation, and phase separation.

Our main results revealed that 14-3-3 β, γ, ε, ζ, and θ can undergo phase separation under *in vitro* molecular crowding conditions as well as heterotypic condensation with α-syn, In the case of amyloid aggregation, the ε, θ isoforms proved to be the most amyloidogenic, forming characteristic short fibrils, which also accelerated α-syn aggregation.

## Methodology

### Multiple Sequence Alignment and Prediction Servers

Sequences of *Homo sapiens* 14-3-3 isoforms (β, γ, ε, ζ, η, θ and σ) were obtained from the Uniprot database [25]. Sequences were aligned using MUSCLE (Multiple Sequence Comparison by Log-Expectation) [26] with default parameters and edited via Jalview [27]. The sequences were imported to PSPHunter [28] or FuzDrop servers and analyzed using default parameters.

### Cloning

The 6xHis-SUMO-14-3-3 β, γ, ε, η, and θ constructs were derived from pGEX-2TKGST-14-3-3 (Addgene no. 13276, 13280, 13279, 13277, 13281) [2]. The 14-3-3 genes and the SUMO-tag were amplified and fused using standard PCR methods. The products were inserted into a pET-28b(-) vector via NdeI and XbaI restriction sites by standard cloning techniques, yielding 14-3-3s constructs fused to an amino-terminal 6xHis-SUMO tag.

pGEX-2TK-14-3-3 beta, gamma, epsilon, eta, tau GST were gifts from Michael Yaffe (Addgene plasmid # 13276, 13280, 13279, 13277, 13281; http://n2t.net/addgene:13276/13280/13279/13277/13281; RRID:Addgene_13276/13280/13279/13277/13281). pET28-6xHix-14-3-3ζ and σ were gifts from Prof. B.M. Burmann. The plasmids and primers used in this study are listed in Table S1.

### Protein Expression and Purification

The purification of 14-3-3 isoforms was done as previously described [23] with minor modifications. In short, plasmid constructs encoding 6xHis-SUMO-tagged 14-3-3 isoforms (β, γ, ε, ζ, η, θ or σ) were transformed into One Shot BL21 Star (DE3) *Escherichia coli* (Thermo Scientific) cells by heat shock at 42 °C for 45 s. Transformed cells were grown in 100 mL of LB medium containing kanamycin (50 μg/mL) at 37 °C and 220 rpm for 16 h. The culture was transferred to 300 mL ZYM-5052 autoinduction medium [29] with kanamycin (50 μg/mL) and incubated at 30 °C and 220 rpm for 16 h. Cells were harvested by centrifugation (3000 g, 20 min, 4 °C). The biomass was resuspended in a buffer (50 mM HEPES and 1.0 M NaCl, pH 7.5), followed by the addition of lysozyme (0.2 mg/ml) and 1 mM phenylmethylsulfonyl fluoride (PMSF). Cells were lysed by sonication (Sonopuls, VS70T probe; Bandelin) for 30 min at 40% amplitude, with a 15 s on/30 s off cycle. The lysate was centrifuged (18 000 g, 30 min, 4 °C). The supernatant was then filtered through a 0.45 μm pore size filter and applied to a gravity column packed with Ni^2+^ Sepharose 6 Fast Flow resin (Cytiva). The column was washed with 50 mM HEPES and 1.0 M NaCl (pH 7.5) buffer, followed by elution with the same buffer solutions containing 0.05 and 0.5 M imidazole. Fractions containing 6xHis-SUMO-tagged 14-3-3 protein were dialyzed (8000 Da MWCO, Biodesign D106) against a 10 mM potassium phosphate buffer (pH 8.0) for 1 h. Home-made Sentrin-specific protease 1 domain (SENP1) was added to cleave the 6xHis-SUMO tag, and the samples were dialyzed for an additional 18 h against a fresh 10 mM potassium phosphate buffer (pH 8.0). Then, the samples were centrifuged (3160 g, 30 min, 4 °C) and filtered through a 0.45 μm pore size filter. Purification using a gravity column was repeated to collect the flow-through fractions. All fractions were checked by SDS-PAGE. Before size exclusion chromatography, 10 mM EDTA and 10 mM dithiothreitol (DTT) were added to all samples. The samples were then concentrated (10 kDa MWCO, Merck) and filtered through a 0.22 μm pore size filter. Size exclusion chromatography was performed using a Tricorn 10/300 (Cytiva) column packed with Superdex 75 resin (Cytiva) and calibrated with Phosphate-Buffered Saline (PBS) (pH 7.4) solution. The collected fractions were checked by SDS-PAGE and concentrated (10 kDa MWCO, Merck). The protein samples were stored at −80 °C.

α-syn and mCherry-α-syn were purified as described previously [30]. During the final purification step involving size-exclusion chromatography, both proteins were exchanged into PBS (pH 7.4). The proteins were then concentrated to 600 µM, distributed into 1.5 mL test tubes (500 µL in each) and stored at -20°C.

### Aggregation kinetics and liquid-liquid phase separation

Samples for the aggregation kinetics assay were prepared by mixing 100 µM of each purified 14-3-3 isoform in PBS (pH 7.4) with 50 µM amyloid-specific Thioflavin-T (ThT) dye. Each sample was divided into four wells of a 96-well non-binding plate, with each well containing a 3 mm glass bead (Merck, cat. No 1040150500). Aggregation kinetics were measured every 10 min at 37 °C with constant shaking at 600 rpm between the measurements using a CLARIOstar Plus microplate reader. ThT dye was excited at 440 nm, and the emission signal was recorded at 480 nm.

For co-aggregation of 14-3-3 and α-syn, 14-3-3ε and 14-3-3θ aggregates were first prepared as described previously and then mixed with α-syn monomer at 2:1 ratio, yielding final concentrations of 100 µM α-syn monomer and 50 µM aggregated 14-3-3. The samples were then further incubated under the same aggregation conditions. For the α-syn seeding experiment, 100 µM of α-syn was first aggregated separately under the same conditions. The resulting aggregates were then further incubated with monomeric α-syn at 2:1 ratio (100 µM of α-syn monomer was mixed with 50 µM α-syn aggregates).

For LLPS studies, samples were prepared identically as for aggregation kinetics, but with the addition of 10% or 20% of the crowding agent polyethylene glycol solution (PEG, 20 kDa, Thermo Scientific), without the 3-mm glass bead and constant shaking. ThT fluorescence and optical density at 600 nm (OD_600_) were measured every 10 minutes at 37 °C. For LLPS studies of 14-3-3 and α-syn, 100 µM of each 14-3-3 isoform was mixed with 100 µM α-syn in the presence of 20% PEG solution.

### Congo red binding analysis

Congo Red (CR) powder was dissolved in 1x PBS (pH 7.4) to a concentration of approximately 5 mM. The solution was then filtered through a 0.22 µm pore-size syringe filter. An aliquot of the solution was diluted 100 times and its absorbance was scanned using a Shimadzu UV-1800 spectrophotometer (ε_486_ = 33 300 M^-1^cm^-1^) in order to determine the exact concentration of the CR stock solution.

100 µM non-aggregated and fibrillar 14-3-3 samples were diluted using 1x PBS and the CR stock solution to a final mixture containing 50 µM 14-3-3 (by monomer concentration) and 10 µM CR. Control samples contained either only 50 µM 14-3-3 or 10 µM CR. After 10 minutes of incubation at room temperature (22°C), the solutions were placed in 3 mm quartz cuvettes, and the sample absorbance was measured between 200 nm and 800 nm using a Shimadzu UV-1800 spectrometer. For each sample, three spectra were averaged and baseline corrected using 1x PBS spectra. In case of fibril samples, the control 14-3-3 sample spectra were subtracted from the spectra of samples with CR.

### Atomic Force Microscopy (AFM)

The freshly cleaved mica was positively charged by applying 50 μL of 0.5% (3-aminopropyl)triethoxysilane (APTES) on the surface and allowing it to functionalize for 5 min. Subsequently, the mica was washed with dH_2_O and dried with airflow, and the procedure was repeated with 50 μL of the protein sample, which was prepared by diluting it 1:9 for 14-3-3s samples and 1:19 for 14-3-3+α-syn samples with PBS. Imaging was performed using a Dimension Icon microscope (Bruker) operating in tapping-in-air mode with aluminium-coated silicon tips (RTESPA-300, Bruker). The images were processed using Gwyddion 2.66 software [26]. Additional images of each sample are located in the supporting information (Fig. S11).

### Fourier Transform Infrared (FTIR) Spectroscopy

Each purified 14-3-3 isoform was diluted to 100 µM in PBS (pH 7.4) and distributed to 1.5 mL test tubes (400 µL final volume, each test tube contained two 3 mm glass beads (Merck, cat. No 1040150500)). The samples were incubated at 37 °C with constant agitation at 600 rpm agitation for 67 h.

The aggregated samples were centrifuged at 16 900 g for 30 min, after which the supernatant was removed and replaced with 400 μL of D_2_O supplemented with 500 mM NaCl (the addition of NaCl generally improves fibril sedimentation) [31]. The centrifugation and resuspension procedures were repeated four times. After the final step, the aggregate pellet was resuspended in 50 μL of D_2_O containing 500 mM NaCl.

FTIR spectra were acquired as described previously [32] using an Invenio S FTIR spectrometer (Bruker), equipped with a liquid nitrogen-cooled mercury-cadmium-telluride detector, at room temperature with constant dry-air purging. For each sample, 256 interferograms with 2 cm^−1^ resolution were recorded and averaged. D_2_O containing 400 mM NaCl and water vapor spectra were subtracted from each sample spectrum, followed by baseline correction and normalization to the same 1595–1700 cm^−1^ wavenumber range. All data processing was performed using GRAMS software.

### Far-UV Circular Dichroism (CD) Spectroscopy

Protein aggregation was performed by incubating samples in 1.5 mL microcentrifuge tubes (Eppendorf) with a 3 mm glass bead at 37 °C for 67 hours with constant agitation at 600 rpm. The samples were then placed in a 0.5 mm quartz cuvette, and CD spectra were measured using a J-815 spectropolarimeter (Jasco). For each sample, spectra were recorded between 190 and 260 nm at 0.1 nm intervals. The data was smoothed using a Savitzky-Golay filter with a 3 nm window. Data analysis was performed using ChiraKit software [27].

### Differential Scanning Fluorimetry (DSF)

Samples for the protein stability assay were prepared by mixing 100 µM of each 14-3-3 isoform in PBS (pH 7.4) with 100 μM 8-anilinonaphthalene-1-sulfonic acid (ANS). The ANS concentration was determined using its extinction coefficient (ε_351 nm_ = 5500 M^−1^ cm^−1^). Protein unfolding was monitored with a Rotor-Gene Q instrument (QIAGEN) using the blue channel (excitation, 365 ± 20 nm; detection, 460 ± 20 nm). The unfolding process was initiated by ramping the temperature from 25 to 99 °C at 1 °C/min increments. Data analysis was performed using MoltenProt software [28].

### Fluorescence Microscopy

14-3-3 isoforms were labelled with fluorescein isothiocyanate (FITC) as described previously [30]. The samples for fluorescence microscopy were prepared by mixing 1 µM FITC-labelled 14-3-3 isoform with 99 µM unlabelled isoform in PBS (pH 7.4). PEG (20 kDa) was added to reach final concentrations of 10% and 20%. Each sample was measured immediately after mixing (0 h) and again after 1 h.

15 μL aliquots of each sample were pipetted onto 1 mm glass slides (Fisher Scientific, cat. No. 11572203), covered with 0.18 mm coverslips (Fisher Scientific, cat. No. 17244914) and imaged immediately after preparation. Short movies were acquired using Olympus IX83 microscope with a 40x objective and fluorescence filter cubes (480 nm excitation and 530 nm emission). Four different images were cropped from each movie and analyzed using Fiji software [33]. Full movies and additional fluorescence microscopy images are available in an online data repository at: doi: 10.5281/zenodo.17709463

For fluorescence microscopy of 14-3-3 and α-syn, 1 µM FITC-labelled 14-3-3 isoform together with 99 µM unlabelled isoform was mixed with 1 µM mCherry-labelled α-syn and 99 µM unlabelled α-syn in PBS supplemented with PEG to a final concentration of 20%. 5 μL aliquots of each sample were transferred onto bright-lined Neubauer counting chambers (Hirschmann). Images were captured from the same field of view using separate fluorescence channels: 480 nm excitation and 530 nm emission for FITC-labelled 14-3-3, and 594 nm excitation and 655 nm emission for mCherry-labelled α-syn. The resulting images were overlaid using Fiji software.

### Droplet statistical analysis

To determine the size of droplets, four fluorescence microscopy images of each 14-3-3 isoform were analyzed. Droplets were segmented using color-based thresholding in ImageJ [29] software, followed by particle analysis. Circular particles (circularity value = 1.0) with a minimum of four pixels in size were included in the droplet area analysis.

## Results

### Amyloid and Phase-Separation Potential of 14-3-3 proteins

Interpretation of amyloidogenic and LLPS properties of 14-3-3 proteins is still at an early stage, indicating a gap in characterising their potential function or disease-related impact. To address this, we initially checked for amyloidogenic regions of 14-3-3 proteins using WALTZ [34] and LLPS propensities with PSPHunter [28] or FuzDrop [35] web servers. The WALTZ algorithm identified that all members have aggregation-prone regions between 170 and 185 amino acids (Fig. 1A), specifically within the sixth α-helix of the proteins. Three members (β, ε, ζ) had an additional 145-153 amino acid region (fifth α-helix), suggesting a structurally inherent susceptibility to misfolding or aggregation. Conversely, all proteins had low probability scores (Fig. S1) for LLPS propensities, with 14-3-3ε and 14-3-3ζ having the highest scores (0.4 and 0.37, respectively) according to PSPHunter, and 14-3-3β, 14-3-3ζ, and 14-3-3σ (0.29, 0.22, 0.21) according to FuzDrop. Both predictors identified different LLPS sequences between family members (Fig. S2), PSPHunter showed the N-terminal as the most likely droplet-forming region. FuzDop additionally indicated a strong probability (>0.6) of C-terminal LLPS, and β, γ, σ protein having extra LLPS regions. These predictive variations stem from differing computational approaches. PSPHunter relies on sequence-pattern recognition derived from experimental datasets, whereas FuzDrop calculates LLPS propensity based on the fundamental biophysics of protein disorder and entropic shifts. Alignment of 14-3-3 sequences (Fig. S2) indicated no overlapping aggregation-prone or LLPS regions; however, in general, amino acid sequence variations at the C-terminal regions of different isoforms were noted as seen previously [36]. Additionally, a high consensus of identified 170-185 aggregation-prone region was observed between isoforms.

**Figure 1.**
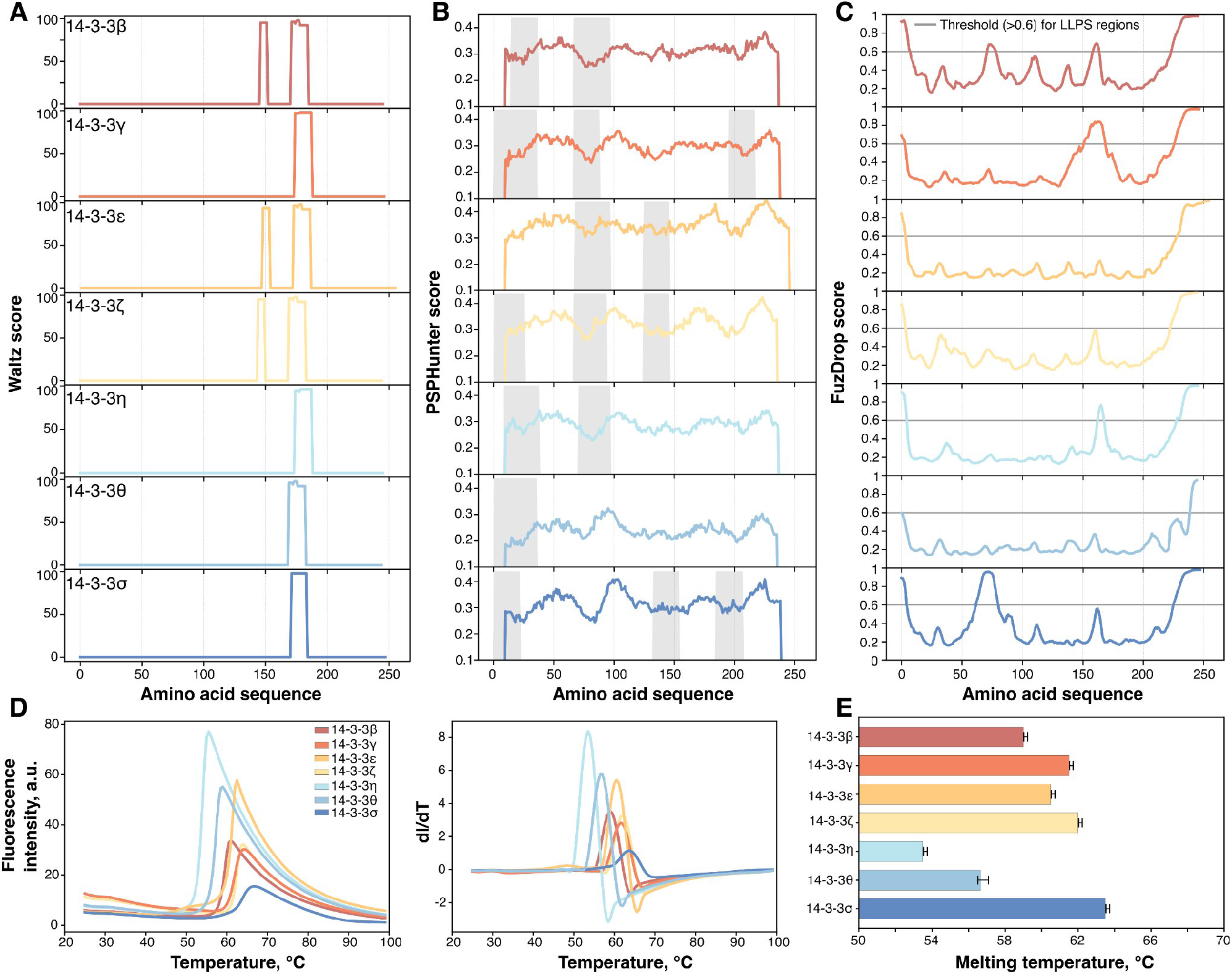
(**A**) Amyloidogenic regions of 14-3-3 isoforms predicted by the WALTZ [34] web server. (**B**) Prediction of LLPS-driving regions by PSPHunter [28] grey areas indicate the most probable phase-separating sequences. (**C**) Prediction of LLPS-driving regions by FuzDrop [35], the grey line indicates the threshold for phase-separating sequences. (**D**) Melting curves, including the first derivatives profiles, of 14-3-3 isoforms and (**E**) calculated melting temperatures by MoltenProt [38]. The error bars represent the standard deviation (n=3).

To supplement the sequence-based aggregation and LLPS propensity, we measured their melting temperature using differential scanning fluorimetry (DSF) to assess their global stability. Between different 14-3-3 members, the unfolding temperature (T_m_) ranged from 53°C to 64°C, with the least stable member being 14-3-3η (53.5 ± 0.1°C) and the most stable -14-3-3σ (63.5 ± 0.1°C). The results closely matched those of previously reported T_m_ values of β, γ, ε, θ, and σ isoforms [37]. The unfolding temperatures of all proteins varied by less than 10°C, indicating similar stability of all isoforms. Overall, 14-3-3 proteins contain aggregation-prone regions but display a low LLPS propensity, with only indications of specific regions with increased LLPS probability. Furthermore, 14-3-3 proteins are thermally stable, as indicated by DSF results. However, due to the limited experimental data available on their amyloid aggregation and phase-separation behaviour, current predictions are highly limited and prone to error as they focus on intrinsically disordered regions or lack information on folded proteins. Hence, we further investigated each propensity experimentally.

### Amyloid aggregation of 14-3-3 proteins

We investigated aggregation properties of each protein using the amyloid specific dye Thioflavin T (ThT) [39], whose fluorescence intensity increases upon binding to fibrils. Aggregation kinetics were monitored over 42 hours at 37 °C, and only the 14-3-3ε sample displayed increased ThT fluorescence, indicating amyloid formation (Fig. 2A). To further characterise amyloid nature of 14-3-3, samples were also mixed with Congo red dye before and after aggregation and their absorbance spectra were recorded (Fig. S3). The spectra of all 14-3-3 aggregates exhibited a characteristic red-shift for amyloid-like structures, with ε, θ, γ and β isoforms showing the largest absorbance. Interestingly, the η isoform bound to CR prior to aggregation and exhibited higher absorbance in its native state compared to after incubation.

**Figure 2.**
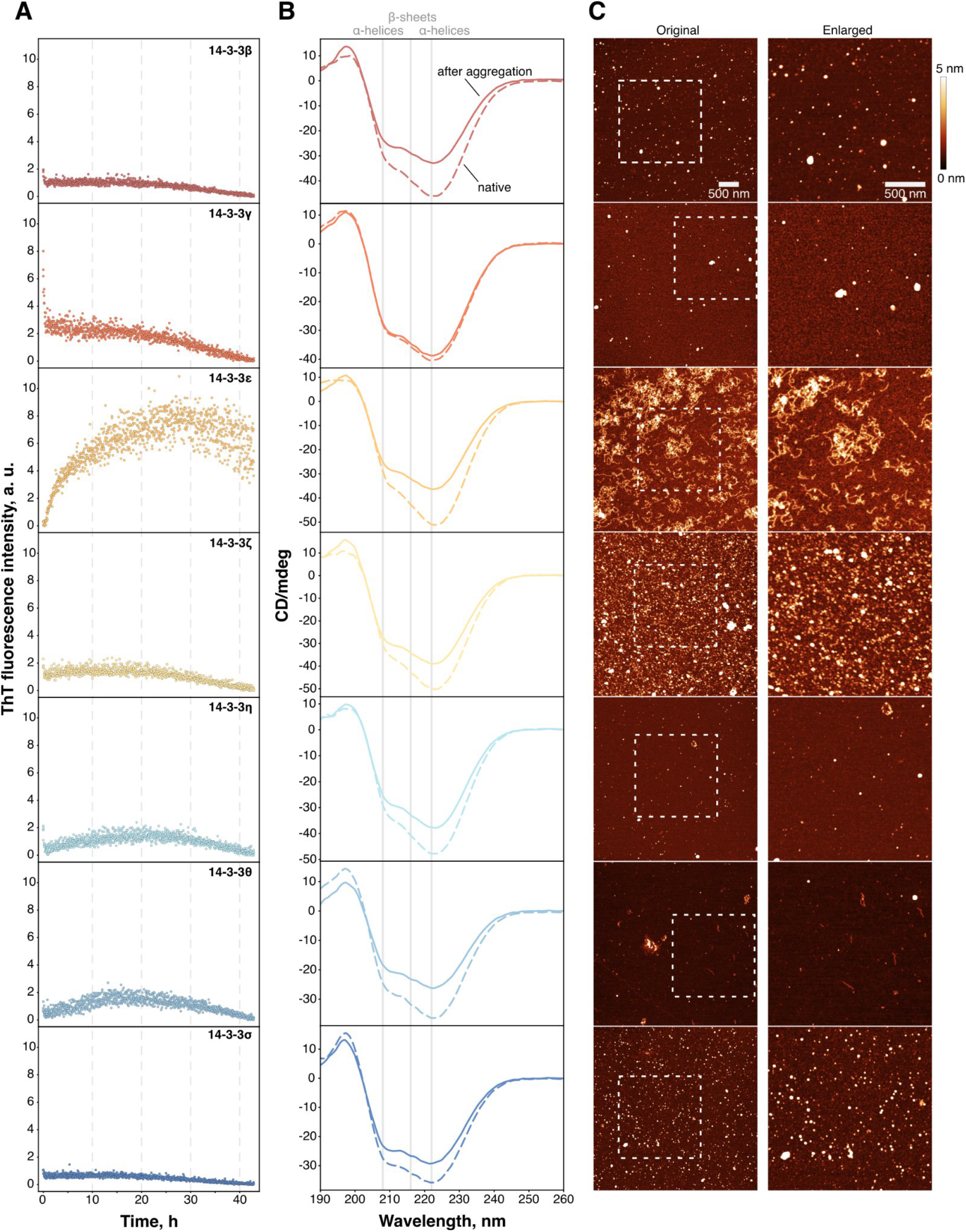
(**A**) Aggregation kinetics of 14-3-3 proteins as observed by ThT fluorescence (**B**). CD spectra of 14-3-3 proteins after aggregation. All samples were incubated for 42 h at 37 °C. The dashed line indicates the native protein spectra, while the solid line represents the spectra after aggregation. (**C**) AFM images of 14-3-3 aggregates (scale bar is 500 nm). The square with dashed lines shows an enlarged part of the image.

Next, we assessed the secondary structure of each isoform before and after 42 h incubation using circular dichroism (CD) (Fig. 2B). As expected, all proteins natively contained α-helices, and after the incubation, the ellipticity increased compared to native spectra, indicating changes in secondary structure or heterogeneity between soluble protein and aggregates. Only 14-3-3γ spectra did not exhibit any changes, which suggests the inherent high stability of the protein. Unfortunately, due to heterogeneous samples, deconvolution of CD spectra was not possible, however to determine if aggregates have β-sheet structures, we isolated them by centrifugation and recorded Fourier transform infrared (FTIR) spectra. β, ε, η, γ isoforms had a sufficient amount of aggregates for detection, and all spectra exhibited a characteristic 1618-1620 cm^-1^ peak (Fig. S4) [40], which indicates the formation of β-sheets, which might stem both from amyloid-like and non-specific aggregates [41]. 14-3-3 ζ, θ, σ aggregates were too small or insufficient to pellet at this stage of incubation, though β-sheet structures in ζ have previously been detected after a longer incubation [23], which was not performed here due to ThT dye bleaching in constant fluorescence measurements [42] and to keep the identical conditions between all isoforms.

Finally, atomic force microscopy (AFM) imaging of each sample showed that most members form unspecific shape aggregates or oligomers, whereas 14-3-3ε and θ produced amyloid-like fibrils with an average height of 1.8 ± 0.4 nm and 1.4 ± 0.3 nm, respectively (Fig. S5), similar to those previously observed in S100A9 [43], prion protein [44] and TDP-43 nucleic acid binding domain [45]. These amyloid-like fibrils typically bind to amyloid dyes, but do not undergo amplified-seed aggregation as long fibrils or do not have a lag phase. However, they could serve as additional surfaces for other protein aggregation or nuclei formation [46].

### LLPS of 14-3-3 proteins

Amyloid aggregation and LLPS often are intertwined processes. The high local concentration of proteins within phase-separated droplets can promote their aggregation [47] and, in some cases, is linked to the initiation of amyloid formation [13]. Currently, there are no data regarding 14-3-3 LLPS behaviour, therefore, we decided to screen them using LLPS-inducing conditions in the presence of PEG.

14-3-3 proteins were labelled with a green fluorescent dye (FITC) and mixed with non-labelled protein in a ratio of 1 to 99, to avoid the influence of FITC modification [30], [48]. The samples for fluorescence microscope imaging were taken immediately and after 1 hour of incubation at room temperature. In the control sample without PEG (Fig. S7), unspecific aggregates or small and isolated droplets were observed. However, at 10% PEG (Fig. S8A) and 20% PEG (Fig. 2A), more evident LLPS and aggregation were observed. The largest condensates were noticed in β, γ, η and θ isoforms, however, across all measured samples, small-sized droplets/aggregates predominated, with a median area ranging from 0.65 to 1.53 µm^2^ (Fig. S9).

To further characterise the process, we also monitored the kinetics of LLPS via turbidity of the solution and ThT fluorescence over 67 h at 37 °C. For most 14-3-3 isoforms, 67 h was insufficient for the signal to reach a plateau, confirming that the transition from LLPS to aggregation of 14-3-3 proteins is a slow process, similar to non-LLPS conditions (Fig. 3B). Only 14-3-3ε aggregation was completed within this period, which coincides with amyloid aggregation. In the presence of 10% PEG, kinetics were also slow, with lower ThT fluorescence intensities (Fig. S7B). Comparison of endpoint values revealed that 14-3-3θ and 14-3-3ε had the highest ThT fluorescence levels, as well as an elevated (>0.5 OD) turbidity, suggesting the formation of larger droplets or aggregation. 14-3-3η initially showed a high OD jump, but decreased during incubation, which corresponds to dissolution of initial aggregates observed in fluorescence microscope images. Droplet fusion was observed in β, γ, ε, ζ, and θ samples (Fig. S10), whereas in others, there were insufficient numbers of droplets or no fusion events. Aggregates were also observed in brightfield images, suggesting that droplets and aggregates are forming at 20% PEG. In comparison to 20% PEG conditions, at 10% of PEG, 14-3-3ε exhibited the highest ThT fluorescence, while 14-3-3θ ThT fluorescence was comparable to other isoforms (Fig. S8C). Furthermore, turbidity was uniformly lower (<0.5 OD) across all samples.

**Figure 3.**
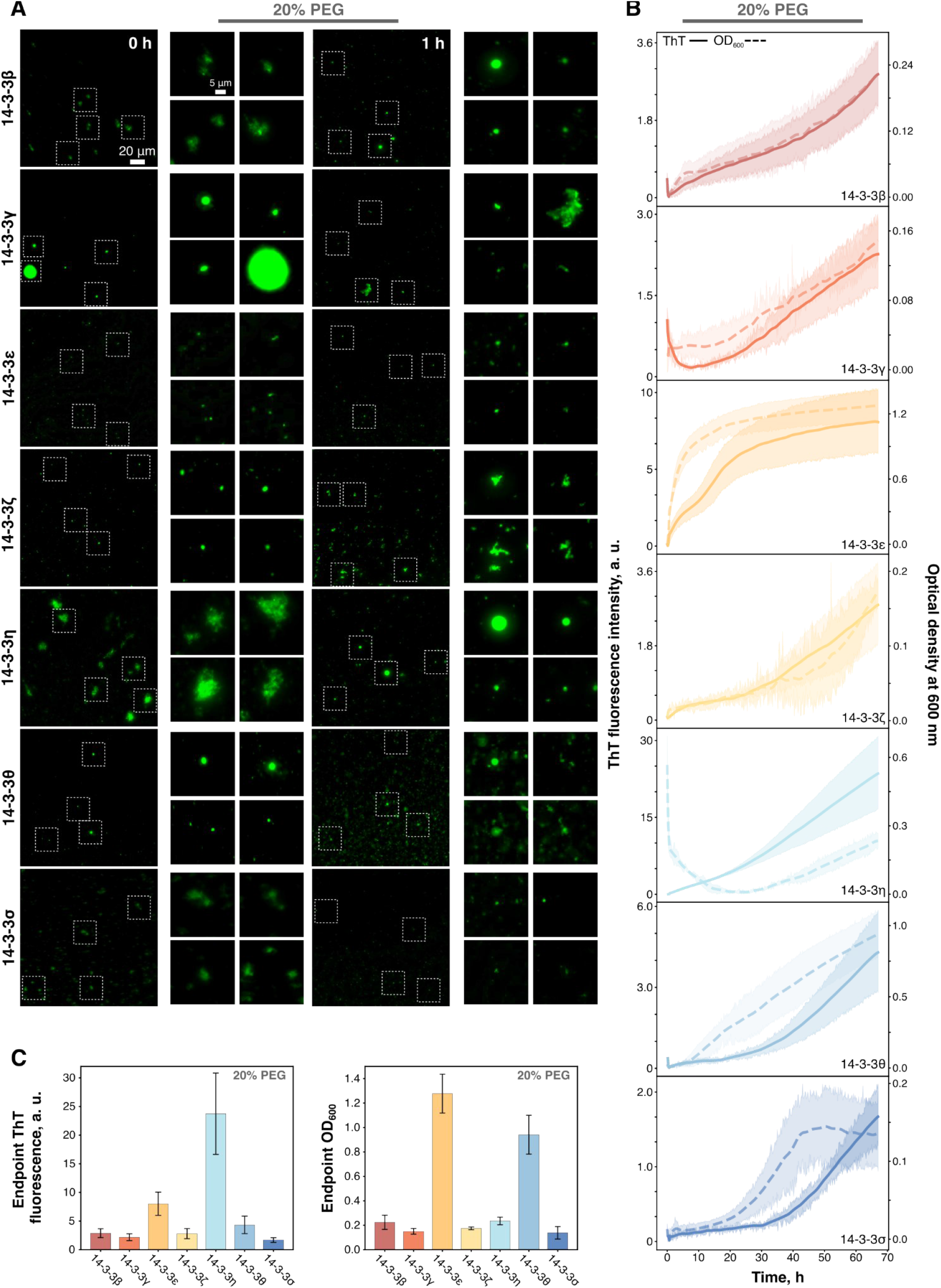
(**A**) Fluorescence microscopy images of 14-3-3 droplets and aggregates formed initially (0 h) and after 1 h (scale bar is 20 µm). Squares with dashed lines indicate enlarged parts of the images (scale bar is 5 µm). (**B**) LLPS kinetics of 14-3-3 proteins in the presence of 20% PEG, followed by ThT fluorescence (solid line) and optical density (dashed line) for 67 h at 37 °C. (**C**) Endpoint (the last measured value) ThT intensities and OD values of LLPS measurements. The error bars represent the standard deviation (n=3).

### 14-3-3 and α-synuclein interaction

14-3-3s are presumed to be potential modulators of LLPS proteins and to interact with them. Particularly, with α-syn, they share even sequence homology [49]. Therefore, we investigated whether 14-3-3 aggregates can accelerate α-syn aggregation or if native proteins can condense into heterotypical droplets. We selected 14-3-3ε and θ isoform aggregates as seeds for α-syn, as amyloid-like filaments were observed within these samples (Fig. 2C). While both isoform aggregates accelerated α-syn aggregation relative to the control, the rate was not as rapid as that induced by α-syn pre-formed aggregates (Fig. 4A). The half time (t_1/2_) value for α-syn control was 21.5 ± 3.4 h, with a lag phase time (t_lag_) of 17.3 ± 2.7 h. In the presence of pre-formed α-syn aggregates, t_1/2_ decreased to 1.2 ± 0.6 h, with no observable lag phase. In comparison, α-syn aggregation induced by pre-formed 14-3-3ε and θ aggregates exhibited similar t_lag_ values of 11.8 ± 0.5 h and 11.2 ± 1.5 h, respectively (Fig. 4B). This suggests that these aggregates likely promote aggregation through secondary pathways, providing a catalytic surface for nuclei formation [46]. Furthermore, the final ThT fluorescence intensities were lower in these samples, which may be a combination of different variables. It could indicate differences in fibril structure, resulting in distinct strains, or the presence of 14-3-3 aggregates/proteins, as it similarly reduced ThT signal intensity during its aggregation. AFM imaging confirmed fibril formation in all samples. The average fibril height for α-syn control was 5.4 ± 0.6 nm, compared to 8.1 ± 1.7 nm in the presence of α-syn seeds. Aggregation induced by pre-formed 14-3-3ε and θ aggregates resulted in lower fibril heights of 4.0 ± 0.4 nm and 3.8 ± 0.5 nm, respectively.

**Figure 4.**
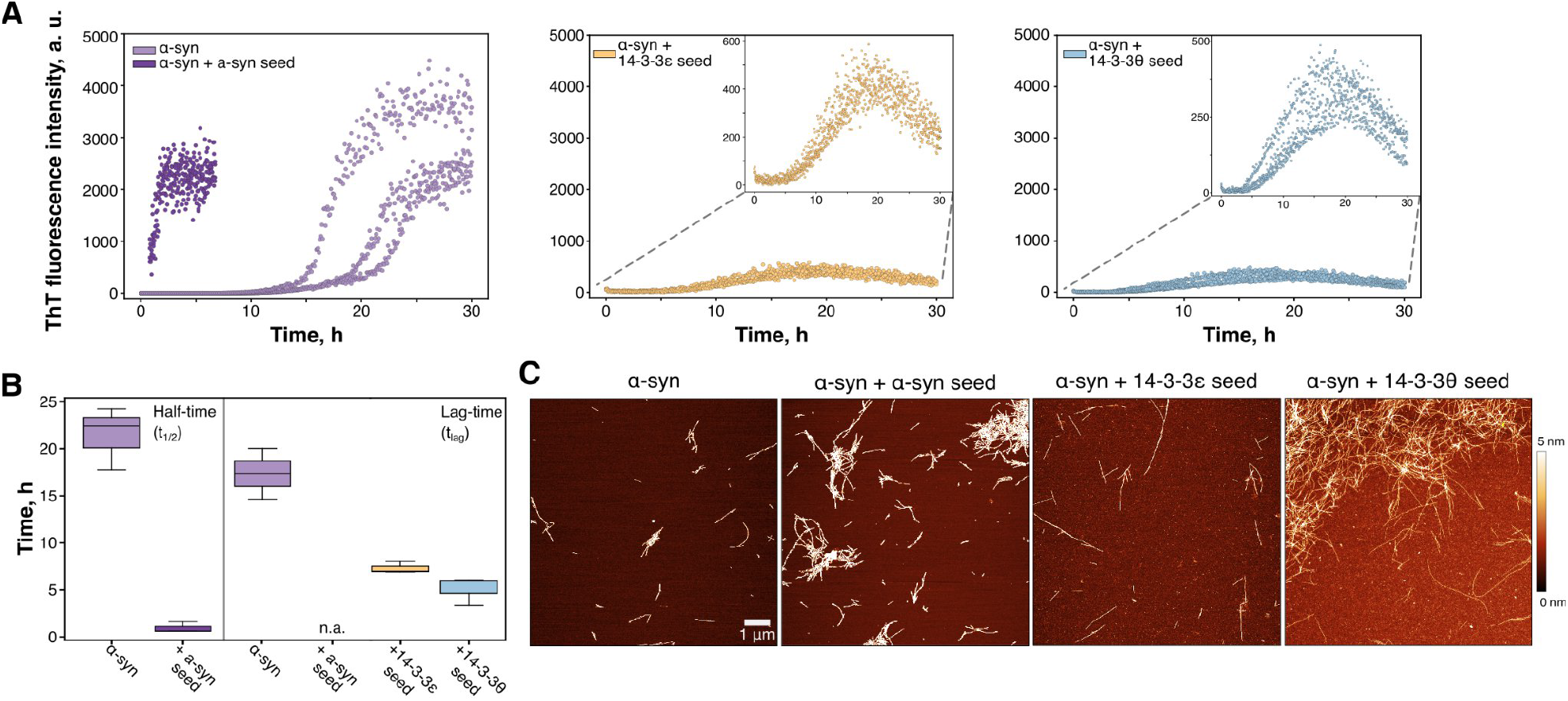
(**A**) ThT fluorescence kinetics of α-syn alone and when seeded with pre-formed aggregates of α-syn, 14-3-3ε and θ (at a 2:1 concentration ratio of α-syn monomer to seed). (**B**) Calculated half-times (t_1/2_) and lag times (t_lag_) derived from the kinetics in (A) with box plots indicating the median, interquartile range (IQR), and whiskers 1.5× range of IQR from box (n=3). (**C**) AFM images of α-syn and co-aggregated samples (scale bar is 500 nm).

We also investigated the condensation of α-syn with 14-3-3 proteins. The LLPS kinetics (Fig. 5A) revealed a stable increase in ThT fluorescence across all samples, with 14-3-3β and η showing the largest values. OD similarly increased in all samples, except when mixed with 14-3-3ε, which resulted in an initial higher OD, followed by a decrease during incubation. Since proteins were mixed at equimolar concentrations (100 µM + 100 µM), we include a sample of α-syn at twice the concentration (200 µM) to compare whether an increase in the total protein concentration or mixing with 14-3-3 samples alters LLPS. The 200 µM α-syn differed from the mixtures; it had the highest OD values, although ThT fluorescence was not as high as with 14-3-3β and η, indicating that 14-3-3 affected the overall dynamics of phase separation. The t_1/2_ values derived from kinetics (Fig. 5B) indicated that the ThT fluorescence increase rate of mixtures was between α-syn 100 and 200 µM samples. The fastest were again 14-3-3β, η (22.1 ± 0.7 and 20.5 ± 0.5 h, respectively), and the slowest was σ (33.7 ± 1.2 h). In general, the addition of 14-3-3 accelerated the formation of ThT-positive aggregates or droplets. Imaging of droplets with a fluorescence microscope confirmed that all 14-3-3s formed heterotypical droplets, except γ. After 1 hour of incubation, 14-3-3γ did not enter α-syn droplets (Fig. S11), whereas other isoforms still showed heterotypical behaviour.

**Figure 5.**
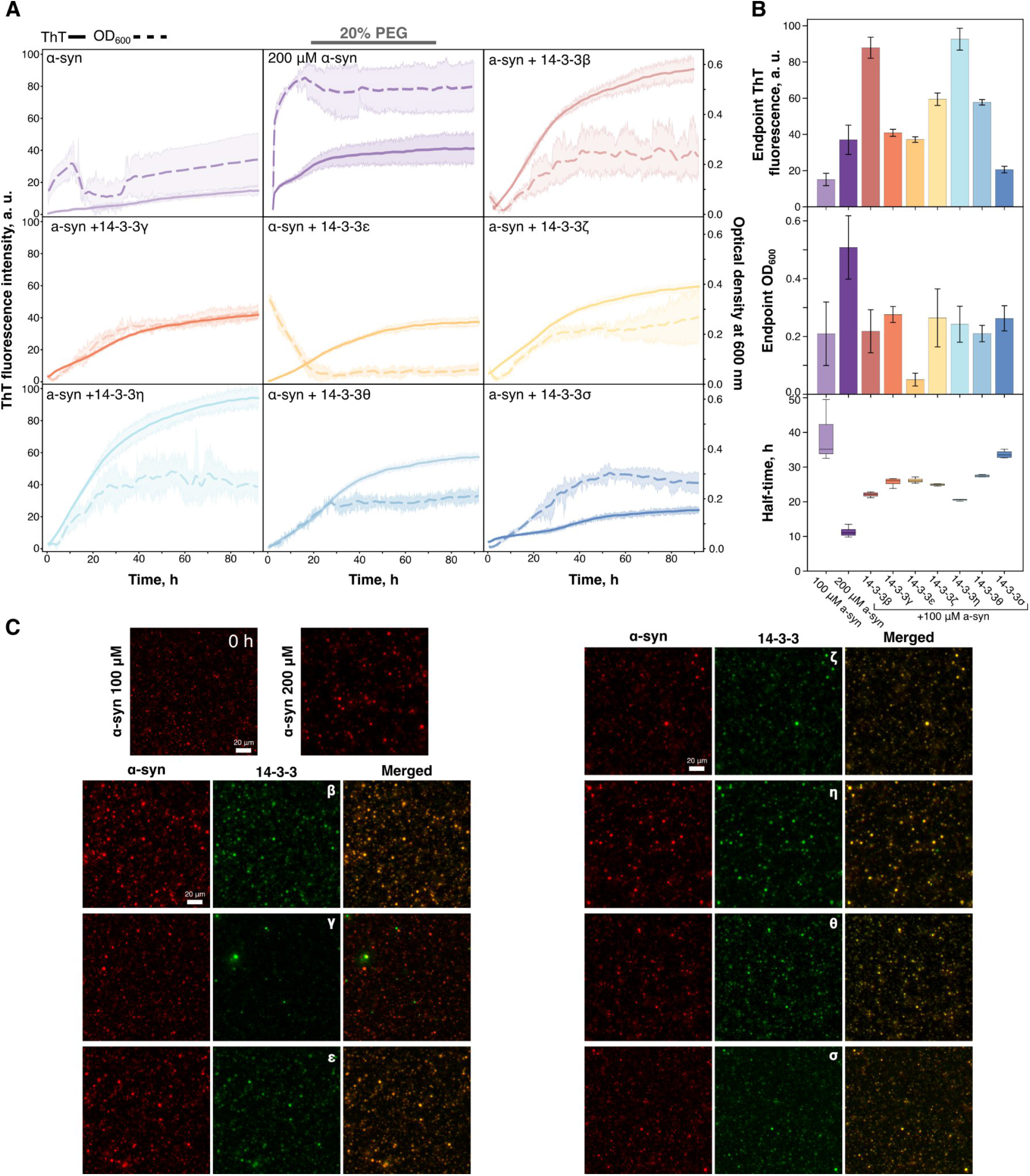
(**A**) LLPS kinetics of α-syn and 14-3-3 proteins in the presence of 20% PEG, followed by ThT fluorescence (solid line) and optical density (dashed line) for 92 h at 37 °C. (**B**) Extracted endpoint fluorescence or optical density and calculated half-times (t_1/2_) derived from the kinetics in (A) with box plots indicating the median, interquartile range (IQR), and whiskers 1.5× range of IQR from box (n=3). (**C**) Fluorescence microscopy images of α-syn (red), 14-3-3 (green) droplets formed initially (0 h) and merged channels (yellow) (scale bar is 20 µm).

## Discussion

The most important function of the 14-3-3 family is acting as a scaffold for various protein interactions. In the cellular environment, interactions between protein-protein or protein-nucleic acids (DNA/RNA) can occur through LLPS, especially in cases with non-globular proteins or proteins found in the nucleus and RNP granules [50]. 14-3-3 can modulate such phase-separated interactions. For example, phosphorylated tau protein undergoes phase separation, but binding to 14-3-3 inhibits LLPS and amyloid aggregation [18]. 14-3-3ζ can also be incorporated within tau droplets and stabilise them, but impede tau droplet-driven tubulin assembly [51]. Similarly, condensates of the human RNA-binding protein SMAUG1 are dissolved by 14-3-3γ or ζ by the co-expression in cells [52]. Another example is LRRK2 kinase, which phosphorylates α-syn, but becomes inactivated upon forming a 14-3-3 complex [5]. Finally, one of the most promoting phase-separating environments in a cell is in the nucleus [53], and 14-3-3 isoforms can be located there. The isoform forming the largest droplets, 14-3-3β, is predominantly localized in neuronal nuclei, whereas 14-3-3γ is found in astrocytic nuclei and 14-3-3ζ in the nuclei of oligodendrocytes [54]. Furthermore, 14-3-3s contain nuclear export signal (NES), which enables them to shuttle other proteins from the nucleus [55], however underlying mechanisms are still not fully understood. With hundreds of known phase-separating partners, it remains unclear how the LLPS of 14-3-3 proteins themselves may influence their binding interactions, as potentially it could create hubs for more efficient binding or prevent unwanted reactions.

Protein aggregation is closely associated with neurodegeneration, both because amyloids can be intrinsically toxic and because aggregation leads to loss of protein function. Recently, elevated levels of 14-3-3γ, β and ζ in cerebrospinal fluid (CSF) have been strongly associated with cognitive decline in early Alzheimer’s disease (AD). Notably, the γ isoform appears to be more prevalent than classical CSF AD biomarkers [56]. Furthermore, loss of the 14-3-3ε gene is associated with increased anxiety, memory dysfunction [57], critical for preventing lissencephaly [58]. Thus, aggregation of 14-3-3ε could potentially contribute to similar dysfunctions. 14-3-3 isoforms ε, θ have already been shown to co-localise with amyotrophic lateral sclerosis (ALS)-linked SOD1 aggregates [59] and ε is a component of prion protein amyloid deposits of Gerstmann-Sträussler-Scheinker disease (GSS) [60]. 14-3-3 isoforms can also recruit proteins in the aggresome formation pathway [61], resolubilize aggregates [62] and act as moonlight chaperones [63]. The concentration of 14-3-3s in the brain is also very high, accounting for up to 1% of all soluble proteins [2], thereby increasing the likelihood of aggregation and LLPS events.

All these studies support the hypothesis that 14-3-3 aggregation may influence, or be influenced by, the aggregation of other proteins. To explore this, we used α-syn as a model client protein, given that its interaction patterns with 14-3-3s have been observed. Under physiological conditions, 14-3-3s can mitigate both α-syn aggregation and fibril-associated toxicity [64], [65]. One of the suggested mechanisms is that 14-3-3 stabilises multimers, allowing α-syn to function, but blocks aggregation pathways [66]. Critically, 14-3-3 also localises within Lewy Bodies [14], placing it directly within the pathological aggregates characteristic of Parkinson’s disease. Our results indicated that 14-3-3 aggregates can accelerate the aggregation of α-syn, or that soluble 14-3-3s can easily incorporate into α-syn droplets during LLPS. These results open a new perspective on how 14-3-3 can interact with its partners, leading to detrimental effects.

## Conclusions

Overall, these results add 14-3-3 to the growing list of potential proteins that can undergo LLPS and form amyloid fibrils *in vitro*. Although many studies have reported their co-localization with aggregates, our work directly demonstrates that 14-3-3 proteins themselves can phase separate into liquid droplets, as well as form heterotypic condensates with α-syn. A limitation of the current study is that we did not investigate these properties *in vivo* or in close to physiological conditions; however given that dysregulated condensate dynamics and protein aggregation are increasingly implicated in neurodegeneration, understanding how 14-3-3 transitions between soluble, condensed, and aggregated states may provide new insights into disease pathogenesis and identify potential therapeutic targets.

## Acknowledgements

We would like to thank Michael Yaffe and Björn M. Burmann for providing the plasmids used in this study.

## Author Contribution

**Saulė Rapalytė: -** Formal Analysis, Investigation, Methodology, Writing - Original Draft, Writing -Review & Editing, Visualization. **Viktorija Karalkevičiūtė -** Formal Analysis, Investigation, Methodology, Writing - Original Draft, Writing -Review & Editing, Visualization. **Ieva Baronaitė**: Formal Analysis, Investigation, Methodology, Writing - Original Draft, Writing -Review & Editing, **Dominykas Veiveris:** Methodology, **Mantas Žiaunys:** Methodology, Writing -Review & Editing **Vytautas Smirnovas**: Resources, Writing -Review & Editing. **Darius Šulskis**: Conceptualization, Formal Analysis, Investigation, Methodology, Writing - Original Draft, Writing -Review & Editing, Visualization.

## Data and materials availability

All data needed to evaluate the conclusions in the paper are present in the paper and/or Supplementary Materials. The raw data used in this paper have been archived and are available on Zenodo: 10.5281/zenodo.17709463. All other relevant data are available from the corresponding author upon reasonable request.

