## Supplementary data for "Liquid-liquid phase separation and amyloid aggregation in the 14-3-3 protein family"

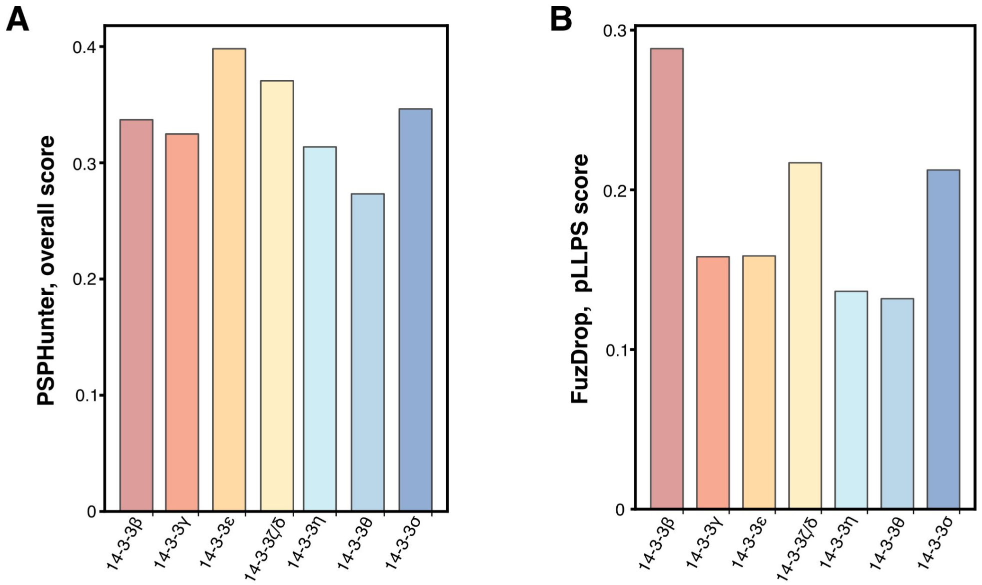


**Figure S1.** PSPHunter (**A**) and FuzDrop (**B**) scores representing predicted phase-separation probability for 14-3-3 isoforms. Lower scores indicate a lower likelihood of phase separation. pLLPS stands for probability of LLPS.

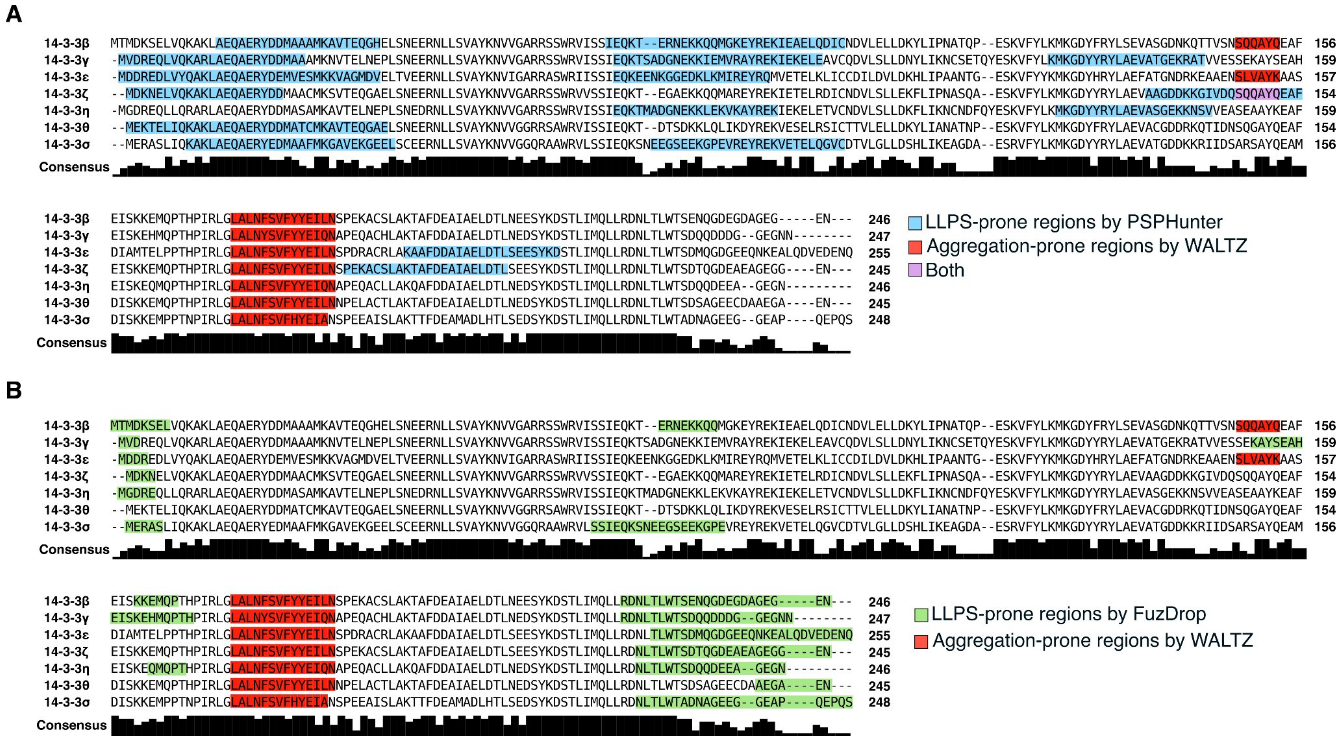


**Figure S2.** Protein sequence alignment of 14-3-3 isoforms by log expectation (MUSCLE). Blue ((**A**) by PSPHunter, green ((**B**) by FuzDrop) colours indicate LLPS-prone regions, red indicates aggregation-prone regions, and purple indicates overlapping regions.

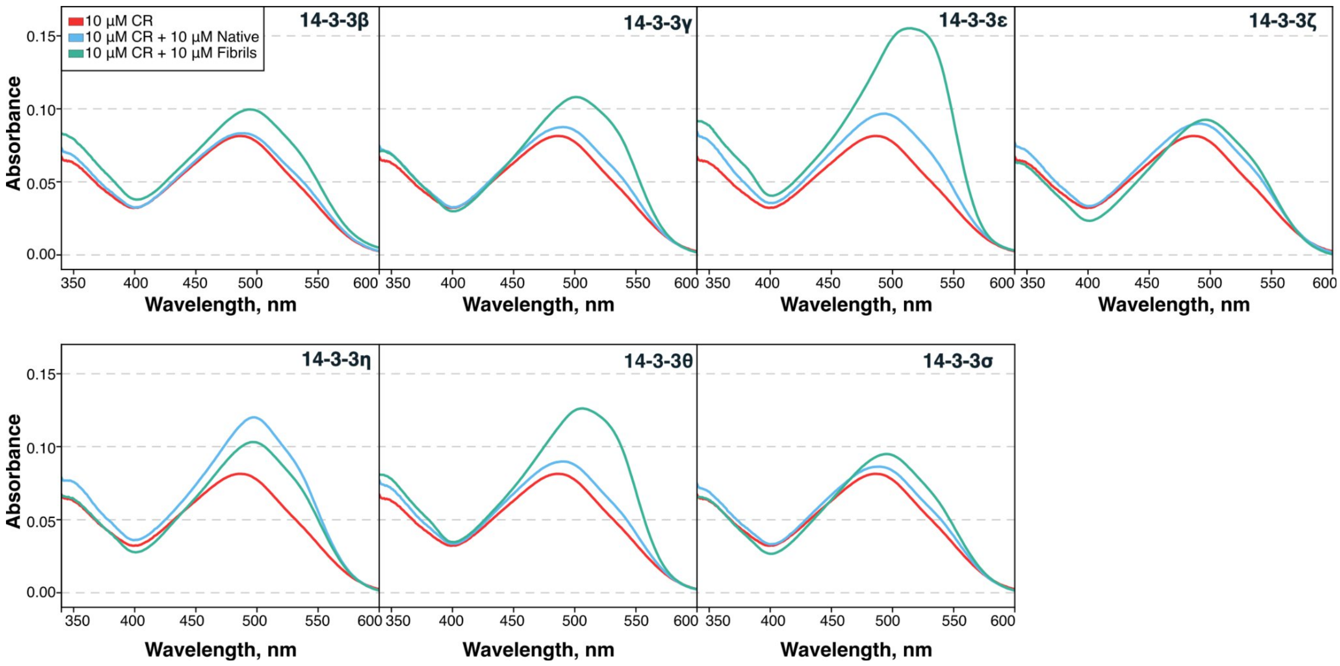


**Figure S3.** Absorbance spectra of Congo red (CR) in the presence and absence of native or aggregated 14-3-3 proteins.

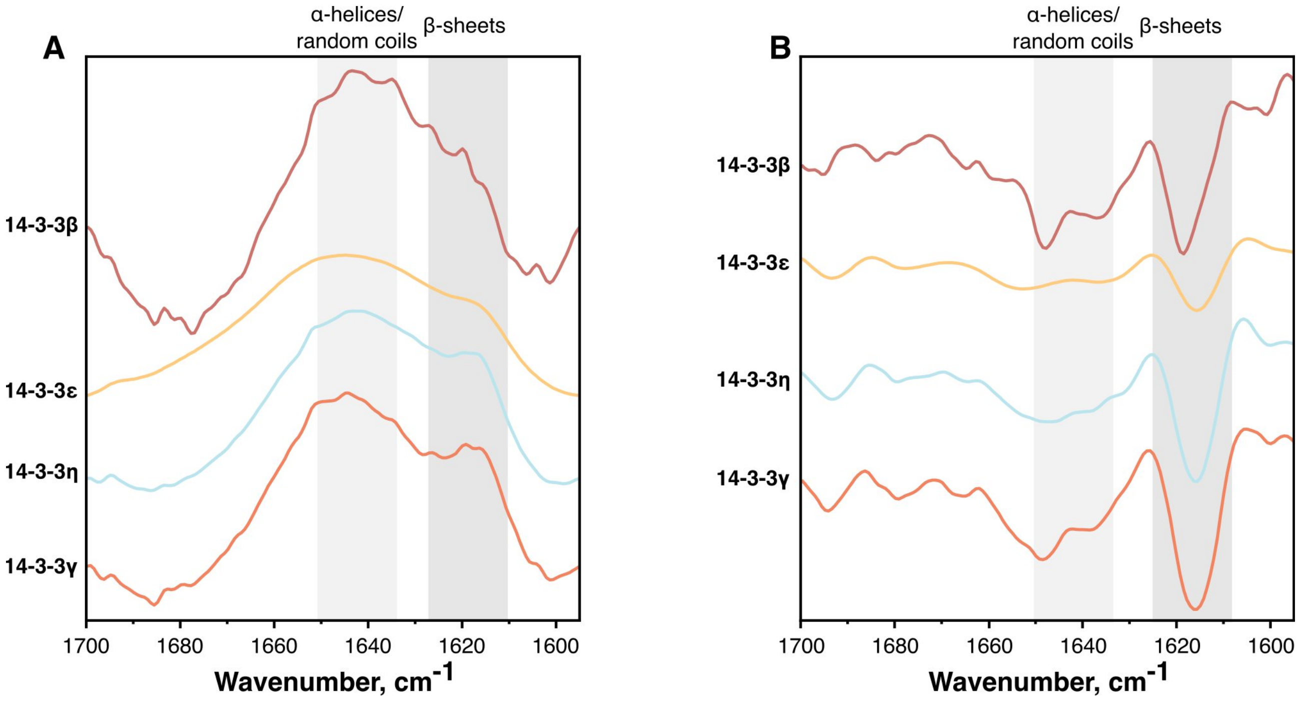


**Figure S4.** FTIR spectra (**A**) and their second derivatives (**B**) of 14-3-3 β, ε, η and γ aggregates.

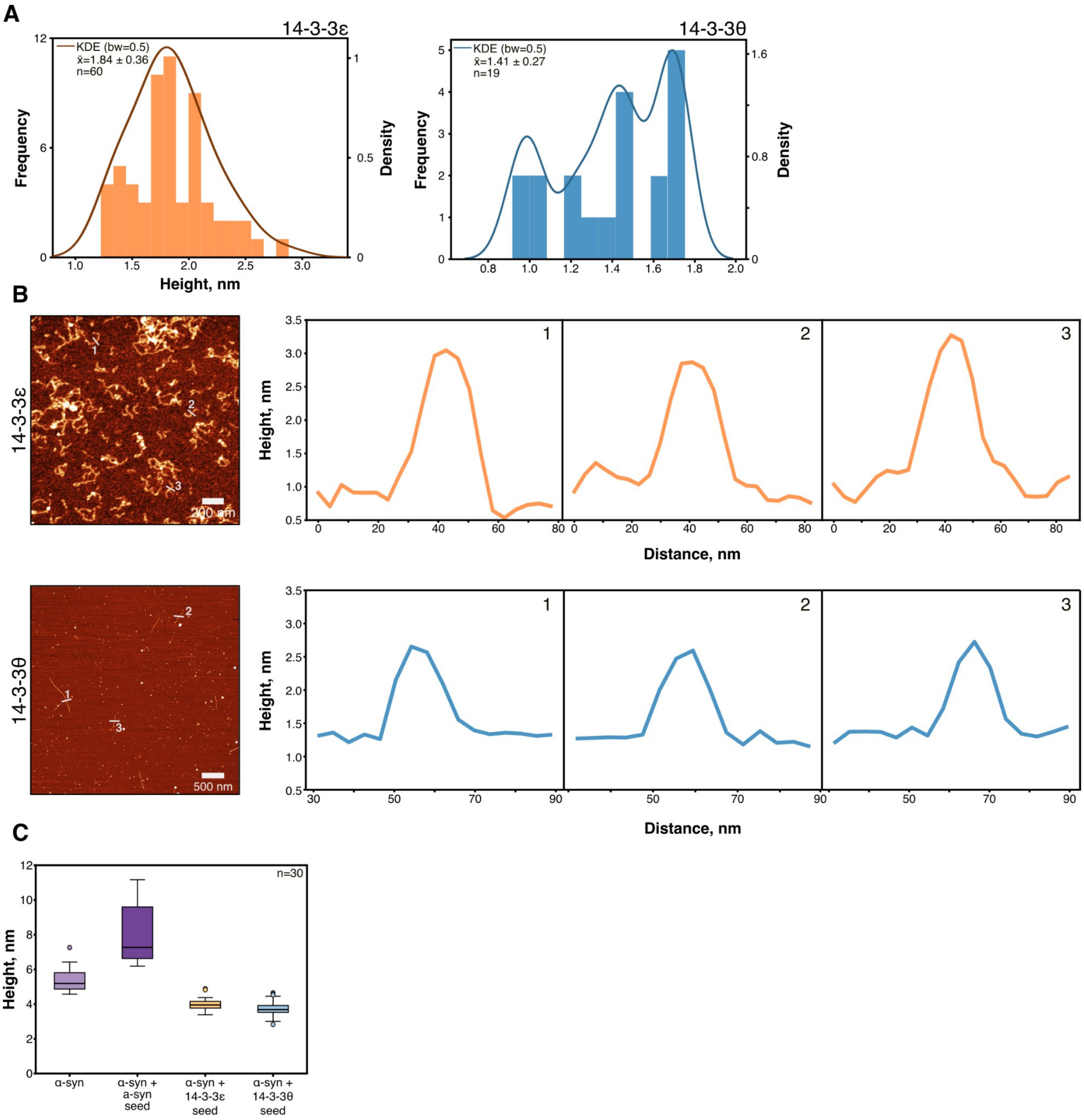


**Figure S5.** (**A**) Height distribution of 14-3-3 ε and θ fibrils. The dark blue line is the kernel density estimation (KDE) function fit. (**B**) AFM images of 14-3-3ε and θ fibrils (scale bars 200 nm and 500 nm, respectively) with selected three profiles of fibril height. (**C**) Comparison of fibril height distributions of α-syn under different seeding conditions. Box plots indicate the median, interquartile range (IQR), and whiskers extending to 1.5× IQR, with outliers shown as individual points (n=30).

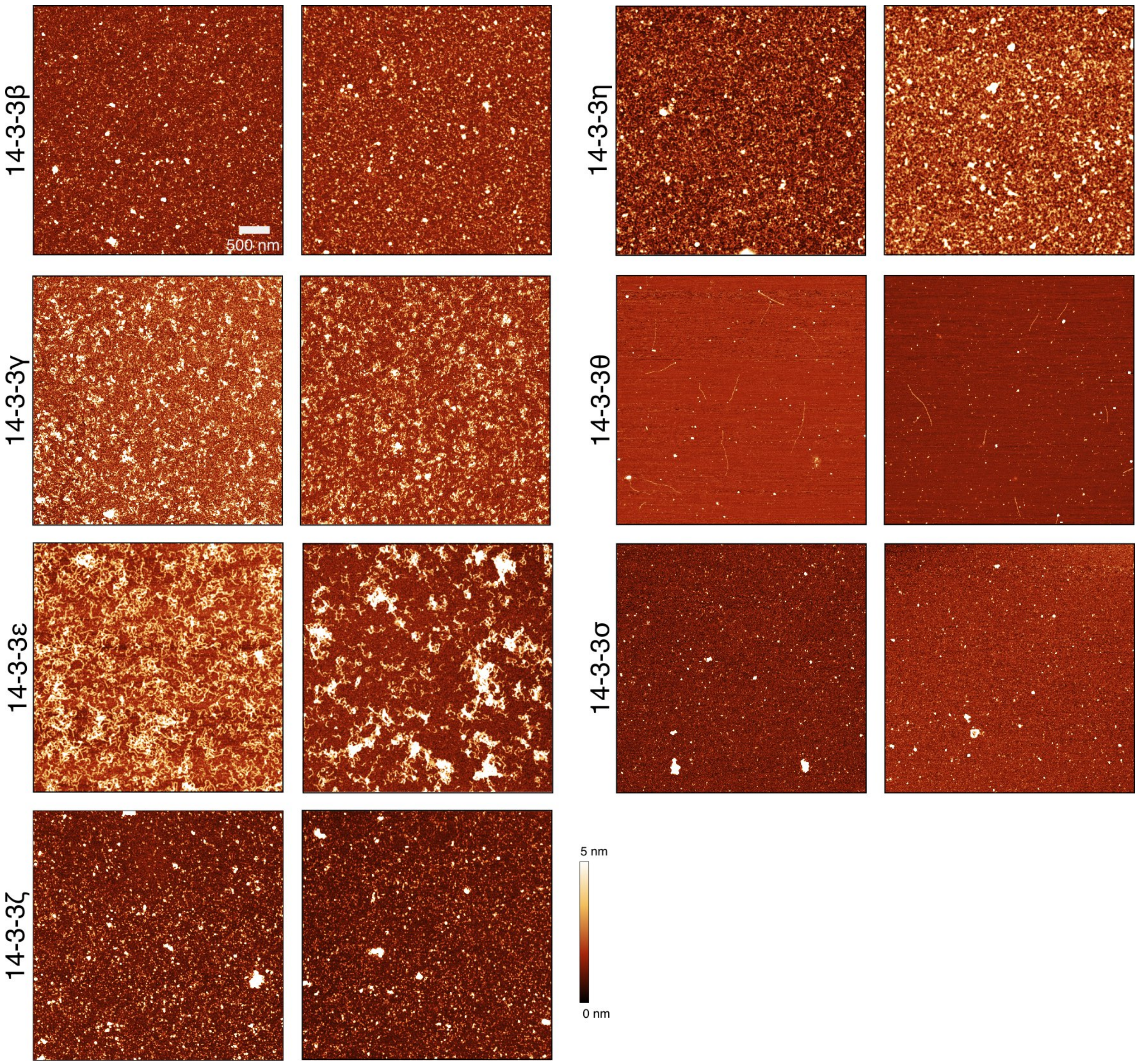


**Figure S6**. Additional AFM images of 14-3-3 aggregates (scale bar is 500 nm).

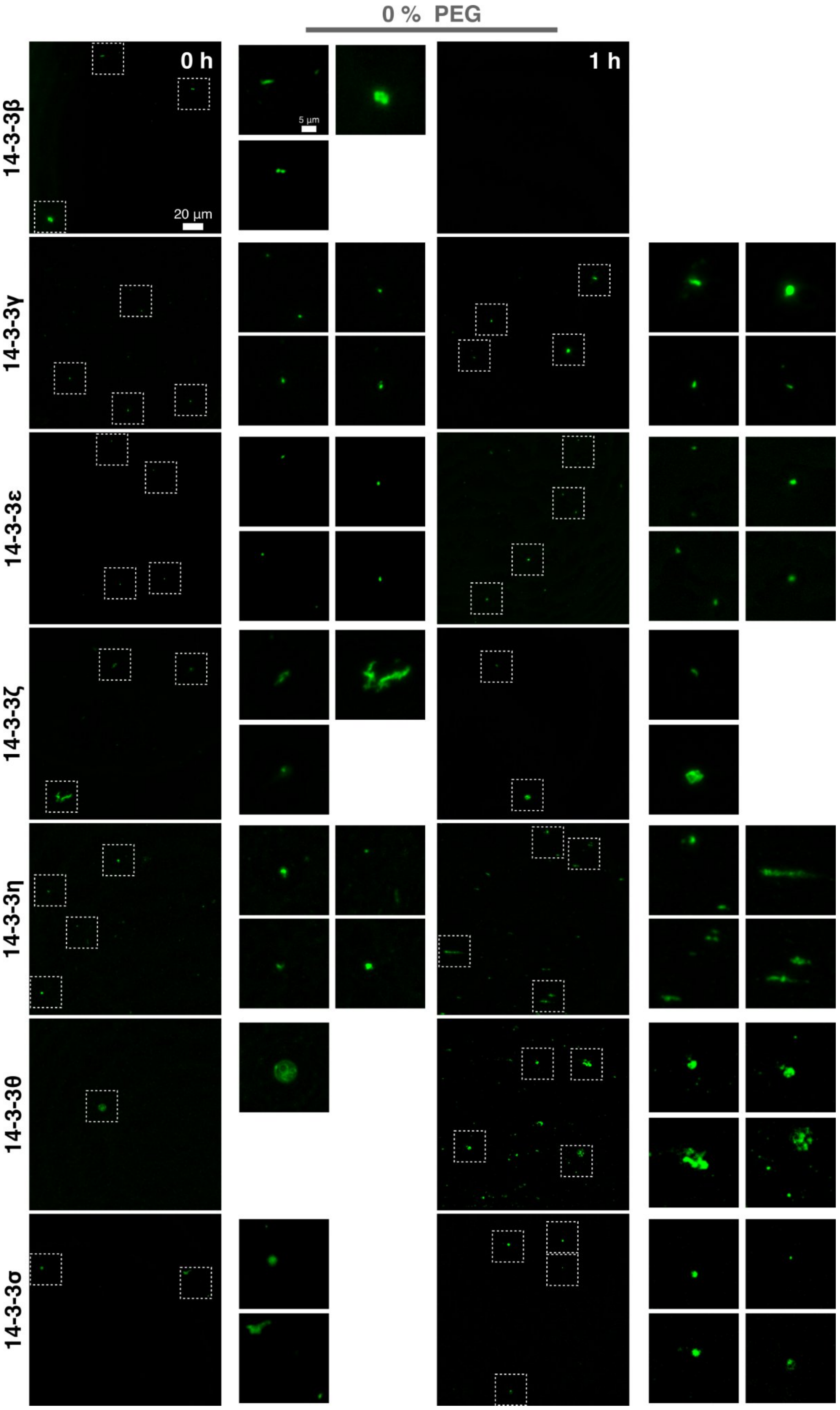


**Figure S7.** Fluorescence microscopy images of 14-3-3 droplets and aggregates formed initially (0 h) and after 1 h (scale bar is 20 µm) without the presence of PEG.

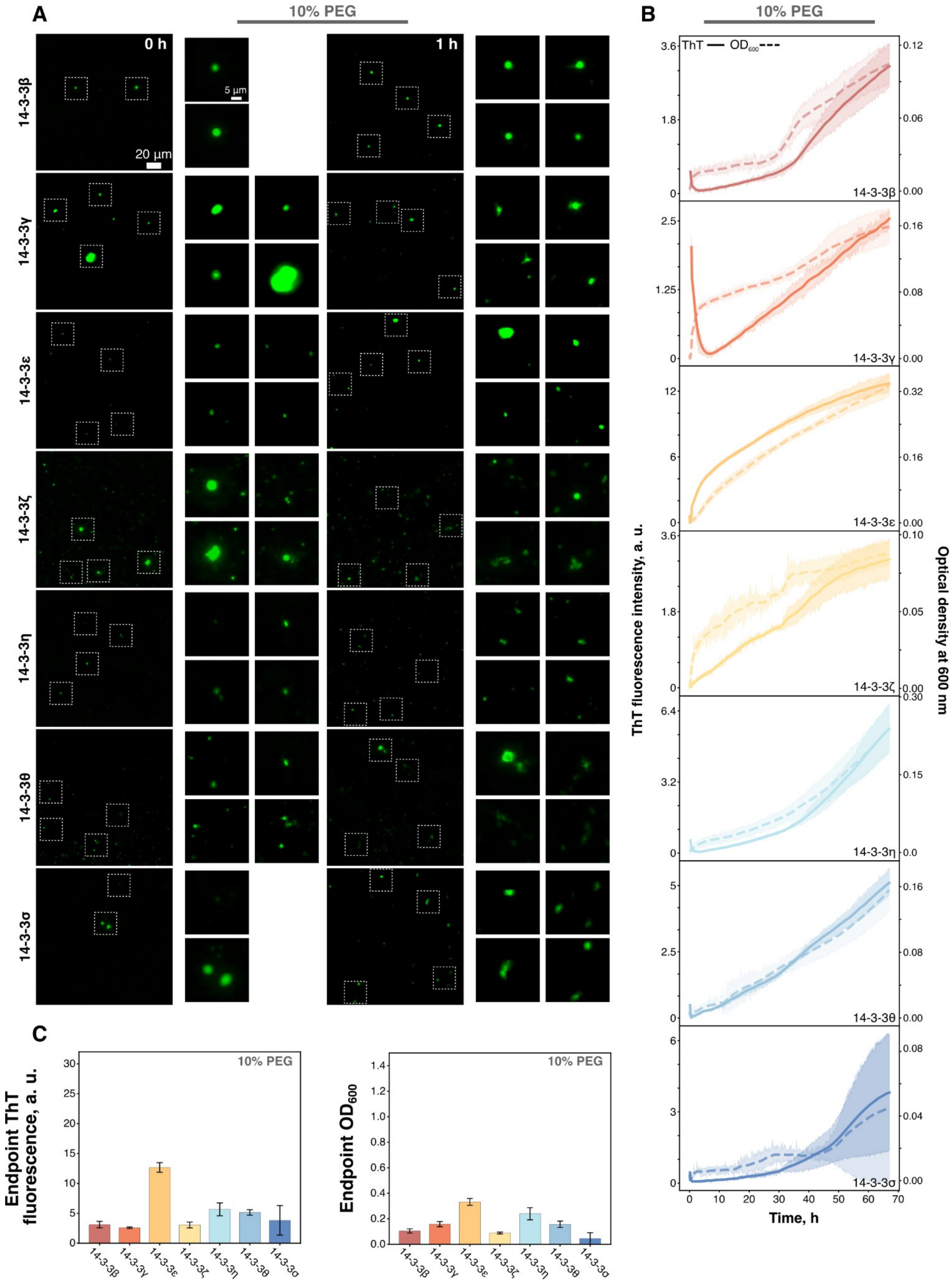


**Figure S8.** (**A**) Fluorescence microscopy images of 14-3-3 droplets and aggregates formed initially (0 h) and after 1 h (scale bar is 20 µm). (**B**) LLPS kinetics of 14-3-3 proteins in the presence of 10% PEG, followed by ThT fluorescence (solid line) and optical density (dashed line) for 67 h at 37 °C. (**C**) Endpoint ThT intensities and OD values of LLPS measurements.

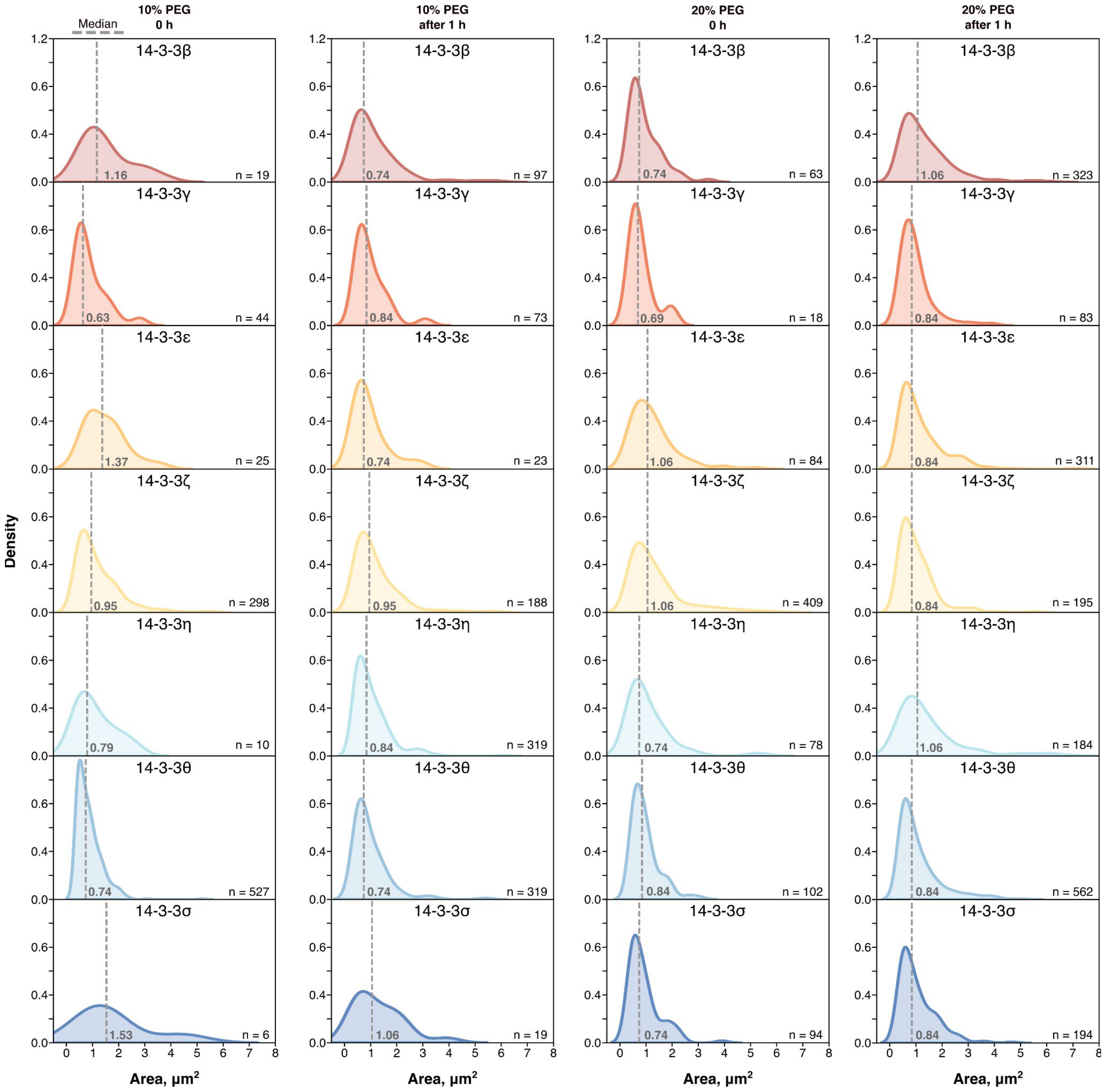


**Figure S9.** Distribution of 14-3-3s droplet size initially and after 1 hour in the presence of 10% or 20% PEG. The dashed line indicates the median.

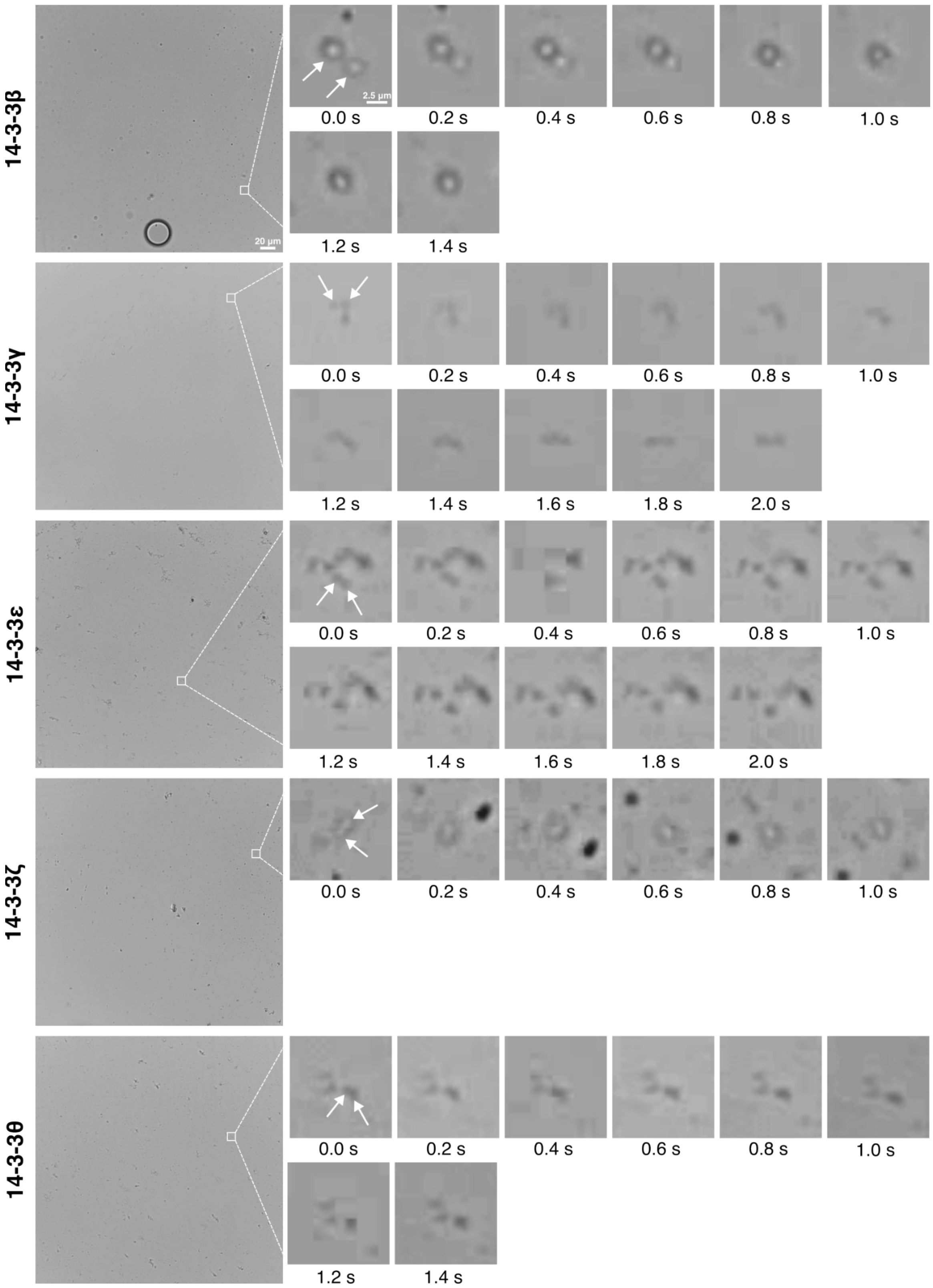


**Figure S10.** Time-lapse microscopy images of 14-3-3s LLPS droplet fusion acquired every 0.2 s (scale bar is 20 µm). White squares indicate enlarged parts of the images (scale bar is 2.5 µm). White arrows indicate fusing droplets.

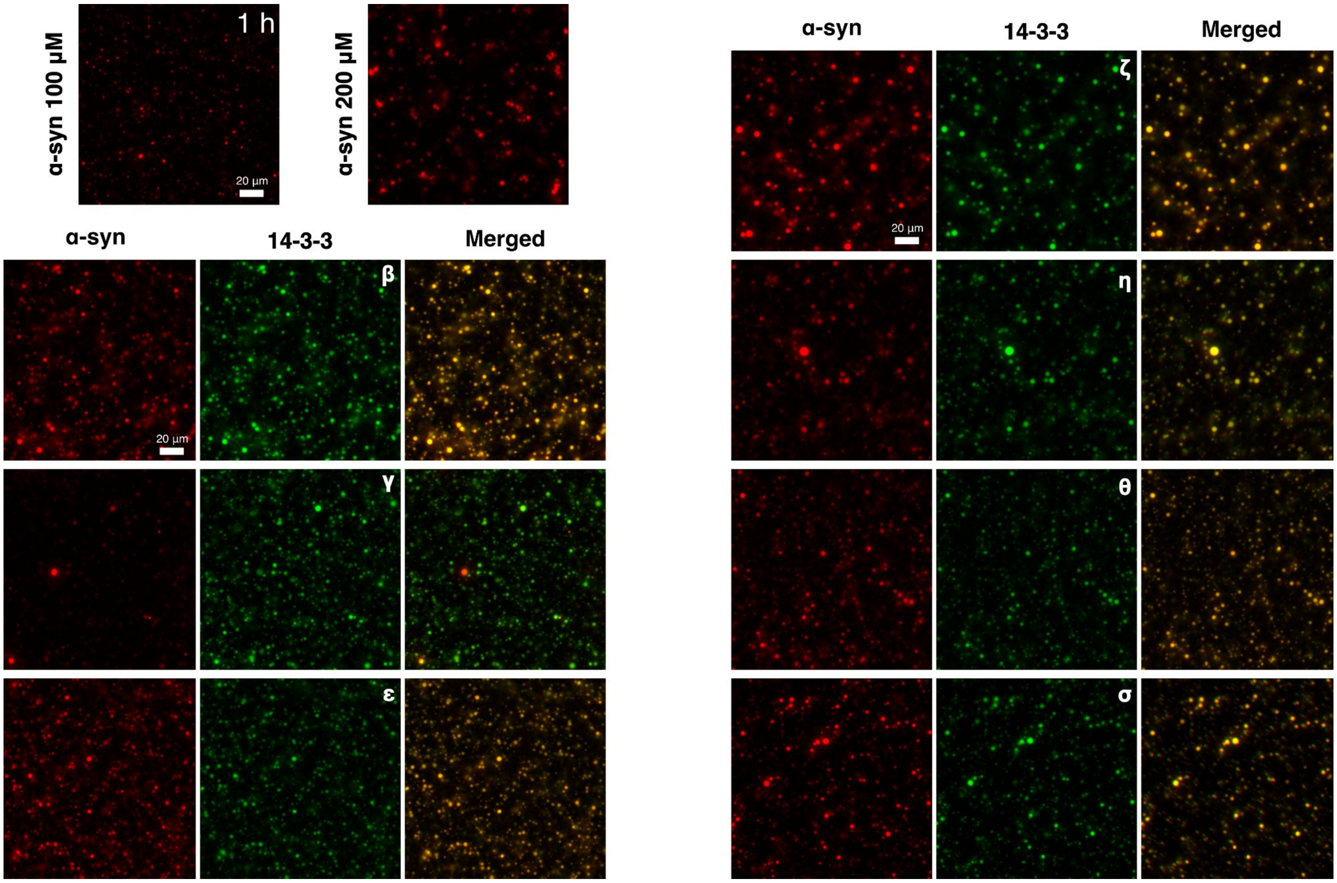


***Figure S11.*** *Fluorescence microscopy images of* α-syn (red), *14-3-3 (green) droplets formed after1h incubation and merged channels (yellow) (scale bar is 20 µm)..*

| Primer | Sequence 5’-3’ |
| --- | --- |
| 14-3-3β_SUMO_end_frw | 5’-GATTGGCGGTatgaccatggacaaaagtgagc-3’ |
| 14-3-3ε_SUMO_end_frw | 5’-GATTGGCGGTatggatgatcgggaggatctgg-3’ |
| 14-3-3η_SUMO_end_frw | 5’-GATTGGCGGTatgggggaccgggagcag-3’ |
| 14-3-3θ_SUMO_end_frw | 5’-GATTGGCGGTatggagaagactgagctgatccag-3’ |
| 14-3-3γ_SUMO_end_frw | 5’-GATTGGCGGTatggtggaccgcgagc-3’ |
| 14-3-3θ_rev | 5’-GCCATATGtcagttttcagccccttctgcc-3’ |
| 14-3-3ε_rev | 5’-GCCATATGtcactgattctcatcttccacatcc-3’ |
| 14-3-3η_rev | 5’-GCCATATGtcagttgccttctcctgcttcttc-3’ |
| 14-3-3γ_rev | 5’-GCCATATGttagttgttgccttcgccg-3’ |
| 14-3-3β_rev | 5’-CGCCATATGttagttctctccctctccagc-3’ |
| SUMO_14-3-3θ_rev | 5’-gtcttctccatACCGCCAATCTGTTCCAGATG-3’ |
| SUMO_14-3-3ε_rev | 5’-gatcatccatACCGCCAATCTGTTCCAGATG-3’ |
| SUMO_14-3-3η_rev | 5’-ggtcccccatACCGCCAATCTGTTCCAGATG-3’ |
| SUMO_14-3-3γ_rev | 5’-ggtccaccatACCGCCAATCTGTTCCAGATG-3’ |
| SUMO_14-3-3β_rev | 5’-ccatggtcatACCGCCAATCTGTTCCAGATG-3’ |
| pET_primer | 5’-ggtcccccatACCGCCAATCTGTTCCAGATG-3’ |
| **Plasmids** | **Description** |
| pET_28_SUMO_14-3-3β | 14-3-3β with fused N-terminal 6xhis-SUMO tag |
| pET_28_SUMO_14-3-3ε | 14-3-3ε with fused N-terminal 6xhis-SUMO tag |
| pET_28_SUMO_14-3-3η | 14-3-3η with fused N-terminal 6xhis-SUMO tag |
| pET_28_SUMO_14-3-3θ | 14-3-3θ with fused N-terminal 6xhis-SUMO tag |
| pET_28_SUMO_14-3-3γ | 14-3-3γ with fused N-terminal 6xhis-SUMO tag |
| pET_28_SUMO_14-3-3σ | 14-3-3σ with fused N-terminal 6xhis-SUMO tag |
| pET_28_SUMO_14-3-3ζ | 14-3-3ζ with fused N-terminal 6xhis-SUMO tag |

**Table 1.** Primers and plasmids used in this study.
